# Establishment of reference standards for multifaceted mosaic variant analysis

**DOI:** 10.1101/2021.08.31.458343

**Authors:** Yoo-Jin Ha, Myung Joon Oh, Junhan Kim, Jisoo Kim, Seungseok Kang, Hyun Seok Kim, Sangwoo Kim

**Author notes:** Corresponding authors: Hyun Seok Kim, Sangwoo Kim. These authors contributed equally to this study (Yoo-Jin Ha, Myung Joon Oh).

## Abstract

Detection of somatic mosaicism in non-proliferative cells is a new challenge in genome research, however, the accuracy of current detection strategies remains uncertain due to the lack of a ground truth. Herein, we sought to present a set of reference standards based on a total of 386,613 mosaic single-nucleotide variants (SNVs) and insertion-deletion mutations (INDELs) with variant allele frequencies (VAFs) ranging from 0.5% to 56%, as well as 35,113,417 non-variant and 19,936 germline variant sites as a negative control. The whole reference standard set mimics the cumulative aspect of mosaic variant acquisition in the early developmental stage owing to the progressive mixing of cell lines with established genotypes, ultimately unveiling 741 possible inter-sample relationships with respect to variant sharing and asymmetry in VAFs. We expect that our reference standards will be essential for optimizing the current use of mosaic variant detection strategies and for developing algorithms to enable future improvements.

## Background & Summary

After conception, postzygotic mutations continuously occur throughout life in humans, causing somatic mosaicism in an individual^1,2^. The variant type, time of origination, and locations of the mosaic mutations result in unique mosaic patterns in a combinatorial manner and further affect phenotypes, including various noncancerous diseases^3–12^. Several efforts have, thus, been made to identify the mutational landscape and mechanisms underlying the mosaic mutations^13–17^.

From a technical aspect, the accurate detection of mutations is at the core of the mosaicism research. To date, conventional bulk sequencing has mainly been exploited by utilizing or modifying variant detection algorithms developed for calling clonal variants, such as cancer mutations^6,18,19^. However, successful application to mosaicism has been obstructed by many challenges, such as low variant allele frequencies (VAF < 10%)^14,17,20,21^ and ambiguity in the use of a control (e.g., variants can exist in control samples)^14,17^. Moreover, fundamentally, there is a severe lack of platforms or materials, known as reference standards, that can be used to measure the detection accuracy of given algorithms, thereby amplifying the confusion regarding the optimal use of tools or algorithms and their reliability. Constructing a standard reference is, thus, a critical first step and serves as the basis for analytical validation and benchmarks for germline and somatic mutations^22–29^. Furthermore, securing a reference standard for mosaic mutations is urgently needed to enable more advanced research.

Herein, we generated robust, large-scale, and cell line-based reference standards using 386,613 single-nucleotide variants (SNVs) and insertion-deletion mutations (INDELs) as positive controls and 35,133,353 negative control positions. The workflow for generating the standard materials and for variant site identification is displayed in Figure 1. The overall idea for the construction aligns with our previous study^30^, as unique germline variants among independent genotypes serve as mosaic variants when mixed in the desired proportions. Initially, six normal cell lines (MRC5, RPE, CCD-18co, HBEC30-KT, THLE-2, and FHC) were prepared and sequenced (1100×) to identify germline variants (see Methods). When MRC5 was employed as an internal reference, each of the five remaining cell lines (RPE, CCD-18co, HBEC30-KT, THLE-2, and FHC) had a unique set of variants, and were called V1 to V5, respectively (Figure 1a; see Table 1 for the full list). When mixed with MRC5 in different proportions, these unique variants are presented as mosaic mutations at designated VAFs.

**Table 1.** Variant set of five cell lines. RPE, CCD-18co, HBEC30-KT, THLE-2, and FHC, represents V1-V5, respectively. Het heterozygous, Hom homozygous

| Variant type | Zygosity | Variant Set |  |  |  |  | Total |
| --- | --- | --- | --- | --- | --- | --- | --- |
|  |  | V1 | V2 | V3 | V4 | V5 |  |
| SNV | Het | 2,698 | 6,158 | 2,058 | 2,127 | 5,825 | 18,866 |
|  | Hom | 133 | 350 | 188 | 129 | 349 | 1,149 |
| INDEL | Het | 89 | 212 | 65 | 60 | 173 | 599 |
|  | Hom | 5 | 12 | 7 | 7 | 16 | 47 |
| Total |  | 2,925 | 6,732 | 2,318 | 2,323 | 6,363 | 20,661 |

**Figure 1.**
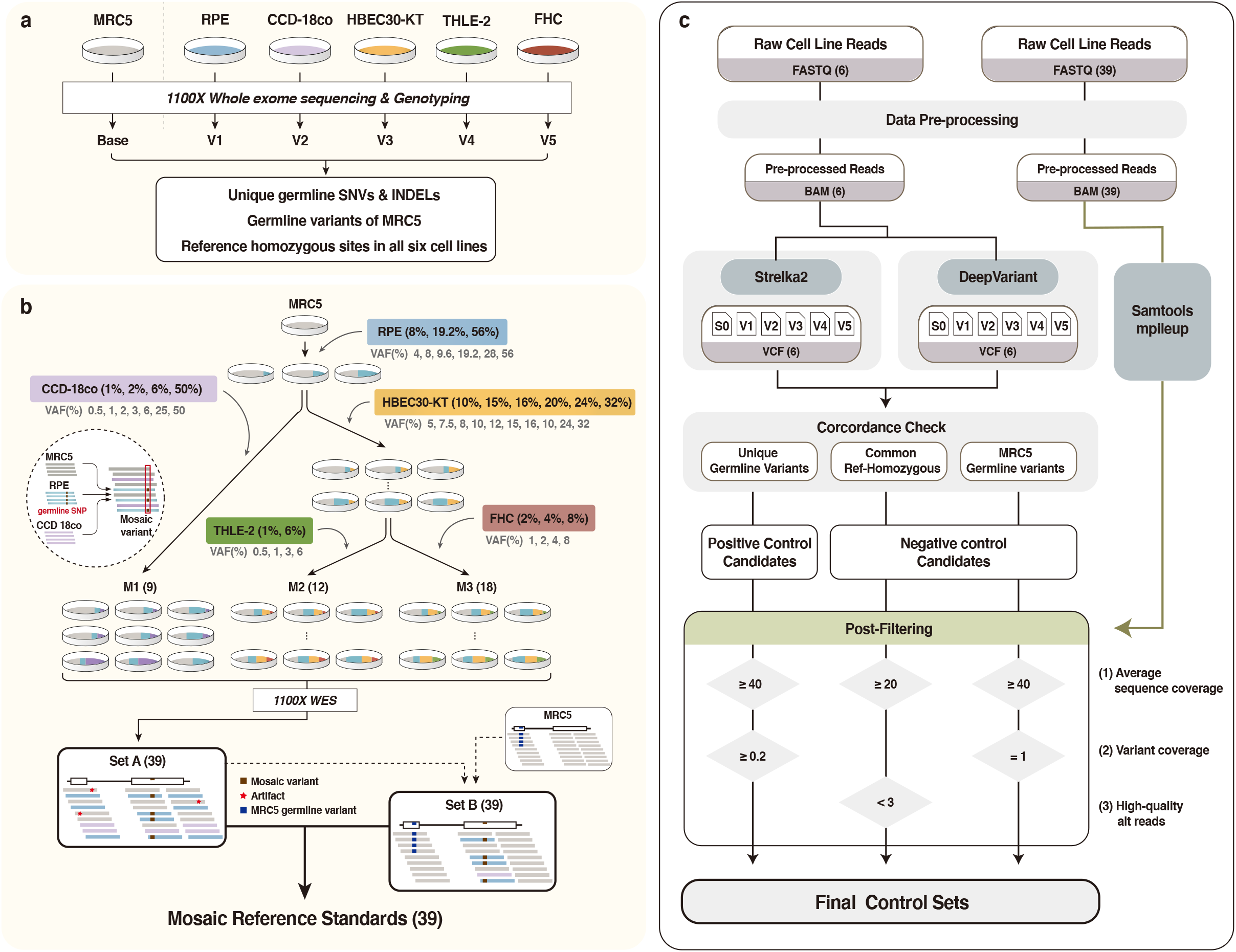
Overall workflow of mosaic reference standard construction. (a) Schematic of the genotyping of six cell lines used as materials. (b) Construction of 39 mosaic reference standards by mixing genetic materials of the six cell lines. Thirty-nine pairs of Set A and Set B were generated by different combinations and proportions of the six cell lines. Set A, sequencing data of the original mixtures; Set B, MRC5 sequencing data with replacement of sequences of variant sites from Set A. (c) Pipeline to generate positive and negative controls in the reference standards. After BAM file preprocessing, candidates for controls were cross-checked using Strelka2 and DeepVariant. Final control sets were fixed with three post-filters using raw read counts (pileup) of 39 mixtures and MRC5 WES data. WES Whole exome sequencing

The mixing procedure was systematically designed to cover a wide range of VAFs and various sharing scenarios (Figure 1b). RPE was mixed into the internal reference (MRC5) at three different ratios (8, 19.2, and 56%) to enable the presentation of the variants in RPE (V1) at six different VAFs (4, 8, 9.6, 19.2, 28, and 56%), depending on the zygosity (hetero- or homozygous). Similarly, CCD18-co and HBEC30-KT were added into the MRC5/RPE mixture at four and six different ratios, respectively. Finally, THLE-2 and FHC were added into the MRC5/RPE/HBEC30 mixture at two and three different ratios, respectively (Figure 1b upper). After the procedure, three final classes of products were generated: M1 (the mixture MRC5/RPE/CCD18-co), M2 (MRC5/RPE/HBEC30-KT/THLE-2), and M3 (MRC5/RPE/HBEC30-KT/FHC). M1 contains the variant sets, V1 and V2; M2 contains V1, V3, and V4; and M3 contains V1, V3, and V5, whose VAFs varied according to the mixing ratios within the class. Of the 12 (3 in RPE × 4 in CCD-18co), 36 (3 in RPE × 6 in HBEC30 × 2 in THLE-2), and 54 (3 in RPE × 6 in HBEC30 × 3 in FHC) possible products in classes M1–M3, 9, 12, and 18 were selected for redundancy and covering efficiency, and subsequently sequenced to ultra-high coverage (1100×) whole-exome sequencing (WES; see Table 1 for the full list). Overall, 9,657, 7,566, and 11,606 positive control variants were included in M1–M3, respectively, with a wide range of VAFs (0.5%–56%), particularly focusing on low frequencies (<10%) (Table 2).

**Table 2.** The compositions and VAFs of variant sets of thirty-nine products. M1, M2, and M3 refer to the three classes depending on the constituent cell lines and 9, 12, 18 products were generated respectively, according to different mixing ratio. V1 RPE, V2 CCD-18co, V3 HBEC30-KT, V4 THLE-2, V5 FHC, VAF variant allele frequency, Het heterozygous, Hom homozygous

| Category | Product | VAF of variant set (Het% / Hom%) |  |  |  |  | # SNV | # INDEL | Total |
| --- | --- | --- | --- | --- | --- | --- | --- | --- | --- |
|  |  | V1 | V2 | V3 | V4 | V5 |  |  |  |
| M1 | M1-1 | 4.0 / 8.0 | 1.0 / 2.0 | - | - |  | 9339 | 318 | 9,657 |
|  | M1-2 | 4.0 / 8.0 | 3.0 / 6.0 | - | - |  | 9339 | 318 | 9,657 |
|  | M1-3 | 4.0 / 8.0 | 25.0 / 50.0 | - | - |  | 9339 | 318 | 9,657 |
|  | M1-4 | 9.6 / 19.2 | 1.0 / 2.0 | - | - |  | 9339 | 318 | 9,657 |
|  | M1-5 | 9.6 / 19.2 | 3.0 / 6.0 | - | - |  | 9339 | 318 | 9,657 |
|  | M1-6 | 9.6 / 19.2 | 25.0 / 50.0 | - | - |  | 9339 | 318 | 9,657 |
|  | M1-7 | 28.0 / 56.0 | 1.0 / 2.0 | - | - |  | 9339 | 318 | 9,657 |
|  | M1-8 | 28.0 / 56.0 | 3.0 / 6.0 | - | - |  | 9339 | 318 | 9,657 |
|  | M1-9 | 4.0 / 8.0 | 0.5 / 1.0 | - | - |  | 9339 | 318 | 9,657 |
| M2 | M2-1 | 4.0 / 8.0 | - | 5.0 / 10.0 | 0.5 / 1.0 |  | 7,333 | 233 | 7,566 |
|  | M2-2 | 4.0 / 8.0 | - | 5.0 / 10.0 | 3.0 / 6.0 |  | 7,333 | 233 | 7,566 |
|  | M2-3 | 4.0 / 8.0 | - | 8.0 / 16.0 | 0.5 / 1.0 |  | 7,333 | 233 | 7,566 |
|  | M2-4 | 4.0 / 8.0 | - | 8.0 / 16.0 | 3.0 / 6.0 |  | 7,333 | 233 | 7,566 |
|  | M2-5 | 9.6 / 19.2 | - | 5.0 / 10.0 | 0.5 / 1.0 |  | 7,333 | 233 | 7,566 |
|  | M2-6 | 9.6 / 19.2 | - | 5.0 / 10.0 | 3.0 / 6.0 |  | 7,333 | 233 | 7,566 |
|  | M2-7 | 9.6 / 19.2 | - | 8.0 / 16.0 | 0.5 / 1.0 |  | 7,333 | 233 | 7,566 |
|  | M2-8 | 9.6 / 19.2 | - | 8.0 / 16.0 | 3.0 / 6.0 |  | 7,333 | 233 | 7,566 |
|  | M2-9 | 28.0 / 56.0 | - | 5.0 / 10.0 | 0.5 / 1.0 |  | 7,333 | 233 | 7,566 |
|  | M2-10 | 28.0 / 56.0 | - | 5.0 / 10.0 | 3.0 / 6.0 |  | 7,333 | 233 | 7,566 |
|  | M2-11 | 28.0 / 56.0 | - | 8.0 / 16.0 | 0.5 / 1.0 |  | 7,333 | 233 | 7,566 |
|  | M2-12 | 28.0 / 56.0 | - | 8.0 / 16.0 | 3.0 / 6.0 |  | 7,333 | 233 | 7,566 |
| M3 | M3-1 | 4.0 / 8.0 |  | 7.5 / 15.0 |  | 1.0 / 2.0 | 11,251 | 355 | 11,606 |
|  | M3-2 | 4.0 / 8.0 |  | 7.5 / 15.0 |  | 2.0 / 4.0 | 11,251 | 355 | 11,606 |
|  | M3-3 | 4.0 / 8.0 |  | 7.5 / 15.0 |  | 4.0 / 8.0 | 11,251 | 355 | 11,606 |
|  | M3-4 | 4.0 / 8.0 |  | 12.0 / 24.0 |  | 1.0 / 2.0 | 11,251 | 355 | 11,606 |
|  | M3-5 | 4.0 / 8.0 |  | 12.0 / 24.0 |  | 2.0 / 4.0 | 11,251 | 355 | 11,606 |
|  | M3-6 | 4.0 / 8.0 |  | 12.0 / 24.0 |  | 4.0 / 8.0 | 11,251 | 355 | 11,606 |
|  | M3-7 | 9.6 / 19.2 |  | 7.5 / 15.0 |  | 1.0 / 2.0 | 11,251 | 355 | 11,606 |
|  | M3-8 | 9.6 / 19.2 |  | 7.5 / 15.0 |  | 2.0 / 4.0 | 11,251 | 355 | 11,606 |
|  | M3-9 | 9.6 / 19.2 |  | 7.5 / 15.0 |  | 4.0 / 8.0 | 11,251 | 355 | 11,606 |
|  | M3-10 | 9.6 / 19.2 |  | 16.0 / 32.0 |  | 1.0 / 2.0 | 11,251 | 355 | 11,606 |
|  | M3-11 | 9.6 / 19.2 |  | 16.0 / 32.0 |  | 2.0 / 4.0 | 11,251 | 355 | 11,606 |
|  | M3-12 | 9.6 / 19.2 |  | 16.0 / 32.0 |  | 4.0 / 8.0 | 11,251 | 355 | 11,606 |
|  | M3-13 | 28.0 / 56.0 |  | 10.0 / 20.0 |  | 1.0 / 2.0 | 11,251 | 355 | 11,606 |
|  | M3-14 | 28.0 / 56.0 |  | 10.0 / 20.0 |  | 2.0 / 4.0 | 11,251 | 355 | 11,606 |
|  | M3-15 | 28.0 / 56.0 |  | 10.0 / 20.0 |  | 4.0 / 8.0 | 11,251 | 355 | 11,606 |
|  | M3-16 | 28.0 / 56.0 |  | 16.0 / 32.0 |  | 1.0 / 2.0 | 11,251 | 355 | 11,606 |
|  | M3-17 | 28.0 / 56.0 |  | 16.0 / 32.0 |  | 2.0 / 4.0 | 11,251 | 355 | 11,606 |
|  | M3-18 | 28.0 / 56.0 |  | 16.0 / 32.0 |  | 4.0 / 8.0 | 11,251 | 355 | 11,606 |
| Total |  |  |  |  |  |  | 374,565 | 12,048 | 386,613 |

Two different types of reference standards are required to enable complete measurement of mosaic detection accuracy, which differ based on the definition of negative controls. Unlike conventional somatic mutations, calling of mosaic variants is susceptible to two different types of errors: (1) calling non-variant sites (e.g., reference allele) and (2) calling germline variants, the latter of which is caused by the unreliability of controls (e.g., variants shared in control samples). Therefore, we provide two different versions of the final sets—set A and set B (Figure 1b lower). Set A is the sequencing data of the original materials, M1–M3, which uses 35,113,417 non-variant sites as negative controls. Set B is processed data, where the sequencing data (BAM) of non-variant sites are replaced by those of the internal reference (MRC5) to contain 19,936 germline variants; this is because the original germline variants of MRC5 are reduced in set A by the mixing procedure. Accordingly, testing should be carried out in both sets. The final list of negative controls is presented in Table 3.

**Table 3.**
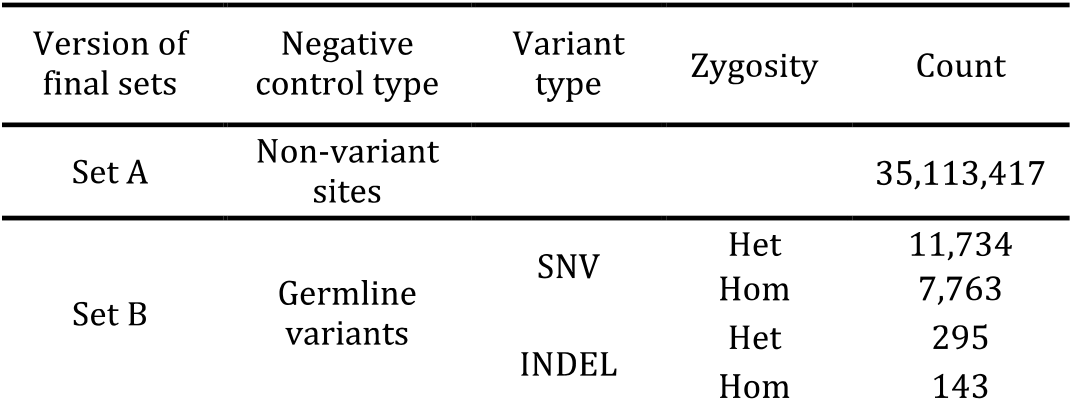
Count of negative controls in final sets. Different types of negative controls are included in the two version of the final sets, Set A and Set B. Het heterozygous, Hom homozygous

Finally, our reference standards allow testing under various realistic biological scenarios by mimicking the structure of multiple lineages in the accumulation of mosaic mutations. There are 741 possible ways to select two samples within M1–M3, each of which provides distinct inter-sample relationships and VAF distributions for mosaic variant detection. For example, M1 and M2 share the variant set V1, whereas V2 is unique in M1, and V3 and V4 are unique in M2. Likewise, M2 and M3 share V1 and V3. In this regard, M2 and M3 are considered closer in the lineage as they have a more recent common ancestor, which can be exploited in more advanced algorithms. The target VAFs display the tendency to decrease in later mutations^1,31,32^. Exceptions caused by the asymmetric doubling of cells and active replication of stem cells or progenitor cells are also considered^3,16^. Owing to these features, our data constitute one of the most comprehensive, versatile, and robust reference standards ever constructed for variant analysis.

## Methods

### Sample collection and preparation

Six normal cell lines (MRC5, RPE, CCD-18co, HBEC30-KT, THLE-2, FHC) were prepared to construct the reference standards. FHC and THLE-2 cells were purchased from the American Type Culture Collection. RPE was purchased from Lonza Bioscience. MRC5 and CCD-18co were purchased from the Korea Cell Line Bank. HBEC30-KT cells were a kind gift from Michael A. White and John D. Minna (University of Texas Southwestern Medical Center). The absence of mycoplasma contamination in all cell lines was verified using the e-Myco VALiD Mycoplasma PCR Detection Kit (LiliF Diagnostics).

All cell lines were cultured in a humidified environment in the presence of 5% CO_2_ at 3°C. FHC cells were grown in DMEM:F12 (Gibco) with 25 mM HEPES (Gibco), 0.005 mg/mL insulin, 0.005 mg/mL transferrin, 100 ng/mL hydrocortisone, 20 ng/mL human recombinant EGF (Thermo Fisher), 10 ng/mL cholera toxin, 10% fetal bovine serum (Gibco), and 1% penicillin–streptomycin (Invitrogen). THLE-2 cells were grown in BEBM (Lonza) supplemented with BEGM Bronchial Epithelial SingleQuots Kit (excluding GA-1000, Lonza), 10% fetal bovine serum, and 1 % penicillin–streptomycin. RPE cells were grown in RtEBM (Lonza) supplemented with RtEGM SingleQuots Supplement Pack (Lonza) and 1 % penicillin– streptomycin. MRC5 cells were grown in MEM (Gibco) with 25 mM HEPES, 25 mM NaHCO_3_, 10% fetal bovine serum, and 1 % penicillin–streptomycin. CCD-18co cells were grown in DMEM with L-glutamine (300 mg/L, Gibco), 25 mM HEPES, 25 mM NaHCO_3_, 10% fetal bovine serum, and 1 % penicillin–streptomycin. HBEC30-KT cells were grown in ACL4 media comprising RPMI 1640 medium supplemented with 0.02 mg/mL insulin, 0.01 mg/mL transferrin, 25 nM sodium selenite, 50 nM hydrocortisone, 10 mM HEPES, 1 ng/mL EGF, 0.01 mM ethanolamine, 0.01 mM O-phosphorylethanolamine, 0.1 nM triiodothyronine, 2 mg/mL BSA, 0.5 mM sodium pyruvate, 2 % fetal bovine serum, and 1 % penicillin–streptomycin.

To achieve the target ratios, mixing was carried out at a DNA level based on the pre-calculated quantities (see Table 2 for final mixture ratios). Genomic DNA was extracted using a QIAamp DNA Mini Kit, according to the manufacturer’s instructions (QIAGEN). A total of 39 mixtures were generated by mixing the genomic DNAs from the six cell lines (see Summary for the procedure). After mixing the genomic DNAs according to the pre-calculated quantities on ice, the mixtures were briefly vortexed, centrifuged, and stored at −20 °C.

### Whole exome sequencing

Exome capture was carried out for six cell lines and 39 mixtures using SureSelect Human All Exon V6 (Agilent Technologies, Inc., CA, USA). To minimize duplicate reads in ultra-deep sequencing, sequencing libraries were constructed two (cell lines) to four (mixture) times for each sample. The quantities of the constructed libraries were evaluated using the 2100 Bioanalyzer Systems (Agilent Technologies, Inc). WES was conducted for the six initial cell lines and 39 mixtures using Illumina NovaSeq 6000 (Theragen Bio Inc.), with targeted read depth of 1100×.

### Processing of the sequencing data

WES reads in FASTQ data were merged and preprocessed using fastp^33^ (0.20.0) to trim overrepresented sequences, such as poly G and adaptors. Reads with low complexity (<30%) were filtered out. The overall sequencing quality was inspected using FastQC (version 0.11.7). All passed reads were aligned to the GRCh38 reference genome using BWA-MEM^34^ (0.7.17). Post-processing, including read group addition, marking PCR duplicates, fixation of mate information, and recalibration of base quality score was applied according to the recommendations of GATK best practices using PICARD (2.23.1) and GATK (4.1.8). We also realigned and left-aligned indels with GATK (3.8.1 and 4.1.5, respectively) to synchronize indel expression in genotyping. Qualimap 2^35^ (2.2.1) was used to calculate the sequencing coverage.

### Genotyping of cell lines

Genotyping of the six cell lines was carried out using two robust germline variant callers: Strelka2^36^ (2.9.10) and DeepVariant^37^ (1.0.0), for autosomal chromosomes, except chr5 [excluded by the copy number variation (CNV) identified in HBEC30, see Technical Validation]. Mutually exclusive SNVs and INDELs (i.e., variants exist in only the cells) were marked as variant sets (V1–V5, see Summary) and were further considered as mosaic variants after mixing.

For SNVs, mutually exclusive variants were collected using the following criteria: (1) variants that were called in both callers and passed the default filtration; (2) variants that were called in only one of the cell lines, with the other five cells being genotyped reference homozygous (i.e., no-call is not allowed); and (3) variants with no signs of copy number alteration (log2 copy number ratio < |0.3| from cnvkit^38^). For indels, similar criteria were applied with an additional rescuing procedure, where single calls (out of two callers) were manually inspected using the Integrative Genomics Viewer^39^ (IGV) for the low concordance among callers^25^. Finally, mutually exclusive variants that passed all criteria in RPE, CCD-18co, HBEC30-KT, THLE-2, and FHC were called V1, V2, V3, V4, and V5, respectively (see Summary). In addition, genotyping of the internal reference (MRC5) was conducted and listed for further processes.

### Finalizing reference standard sets

Genotypes of the 39 mixtures (within M1, M2, and M3) were theoretically pre-fixed by the genotypes of the six cell lines and their mixture compositions. To finalize the reference standard sets, we conducted a series of post-filtration procedures to remove sites that significantly deviated from the expected coverage and VAFs, particularly from extrinsic and systematic errors. The procedures were applied to two difference sets: set A and set B (see Summary) (Figure 1c).

#### Reference standard with non-variant sites as the negative control (set A)

Set A is basically the sequencing data of the 39 mixtures themselves with reference homozygous sites as negative controls that are identified from the genotyping of the six cell lines. Therefore, the finalization of set A only required a few additional filtration steps.

Preprocessed sequencing data were used for the final confirmation of control positive sites based on two filtration criteria: (1) sequencing coverage and (2) variant coverage. Regarding sequencing coverage, raw allele counts were calculated in all targeted positions using SAMtools^40^ mplileup (1.10), ignoring soft or hard clips. For each variant site, the mean coverage of the 39 samples was calculated, and low coverage sites (<40×) were removed; these sites should theoretically be variant positions but cannot be used as positive controls because of the low-sequencing coverage. Likewise, non-variant (negative control) sites with an average coverage of < 20× were removed. Regarding variant coverage, for each variant *v*, variant coverage was defined as (number of samples that actually harbored *v*) / (number of samples designed to harbor *v*). Variants with low variant coverage (<20%) were considered to be affected by low-sequencing efficiency and were, thus, removed. Moreover, non-variant positions with more than three high-quality (BQ ≥ 30) alternative alleles were filtered out to prevent any interference from experimental or systematic bias, rather than sequencing artifacts.

#### Reference standard with germline variants as the negative controls (set B)

Unlike set A, set B requires an additional process to replace germline variant sites of mixtures with those of internal reference (MRC5). Sequences of MRC5 was first down sampled to 1,100× with random seed for 39 times using PICARD DownsampleSam (2.23.1). Reads in the target sites were extracted using bedtools^41^ (2.28.0) and merged into each down-sampled MRC5 data. Before the replacement, we verified that the sequenced fragment length, GC content, and quality of bases were comparable for the two types of data, WES reads of MRC5, and 39 mixtures. Consequently, mosaic variants and germline variants of MRC5 coexisted within set B with the replacement.

A similar post-filtration performed for set A was applied to set B. First, sequencing coverage filtration was equally applied. Second, the VAF in each germline variant site was assessed to filter out sites that violate beta-binomial distribution for heterozygote [74, 76 for α, β calculated from MRC5 heterozygous single-nucleotide polymorphisms (SNPs), two tailed p-value < 0.01] and homozygote (VAF < 0.9) to consider over-dispersion and capture bias in WES. Lastly, variant coverage was calculated to remove germline variants that were missing in any mixture samples (variant coverage < 1).

### Data Records

The raw WES FASTQ files of 6 cell lines and 39 mixtures are available from the Sequence Read Archive under the accession code [PRJNA758606]. Thirty-nine pairs of set A and set B are also available in BAM file format to be readily applied for evaluation of methods. Positive and negative controls of mosaic reference standards are available in GitHub^42^ and Zenodo^43^. The expected VAFs and compositions of positive controls in each sample are presented in Table 2.

### Technical Validation

#### Validation of normal cell line stability

We used six normal immortalized cell lines for stability and reproducibility, as they do not continuously acquire small and large variants during cell culture, unlike cancer cell lines. The distribution of heterozygous SNPs detected using Strelka2 annotated with gnomAD (v2.1.1) showed a singular peak at VAF 0.5 in all six cell lines, demonstrating the monoclonality of the materials (Figure 2a). As positive controls were constructed by mixing independent cell lines, it was important to validate their diploid genotypes. Therefore, the overall regions of all six cell lines appeared to be copy number neutral, except the sex chromosomes and entire chromosome 5 of HBEC30-KT, as commonly observed^44^ (Figure 2b). The unique germline variants used for the positive control were selected from copy number neutral regions through CNV analysis (Methods).

**Figure 2.**
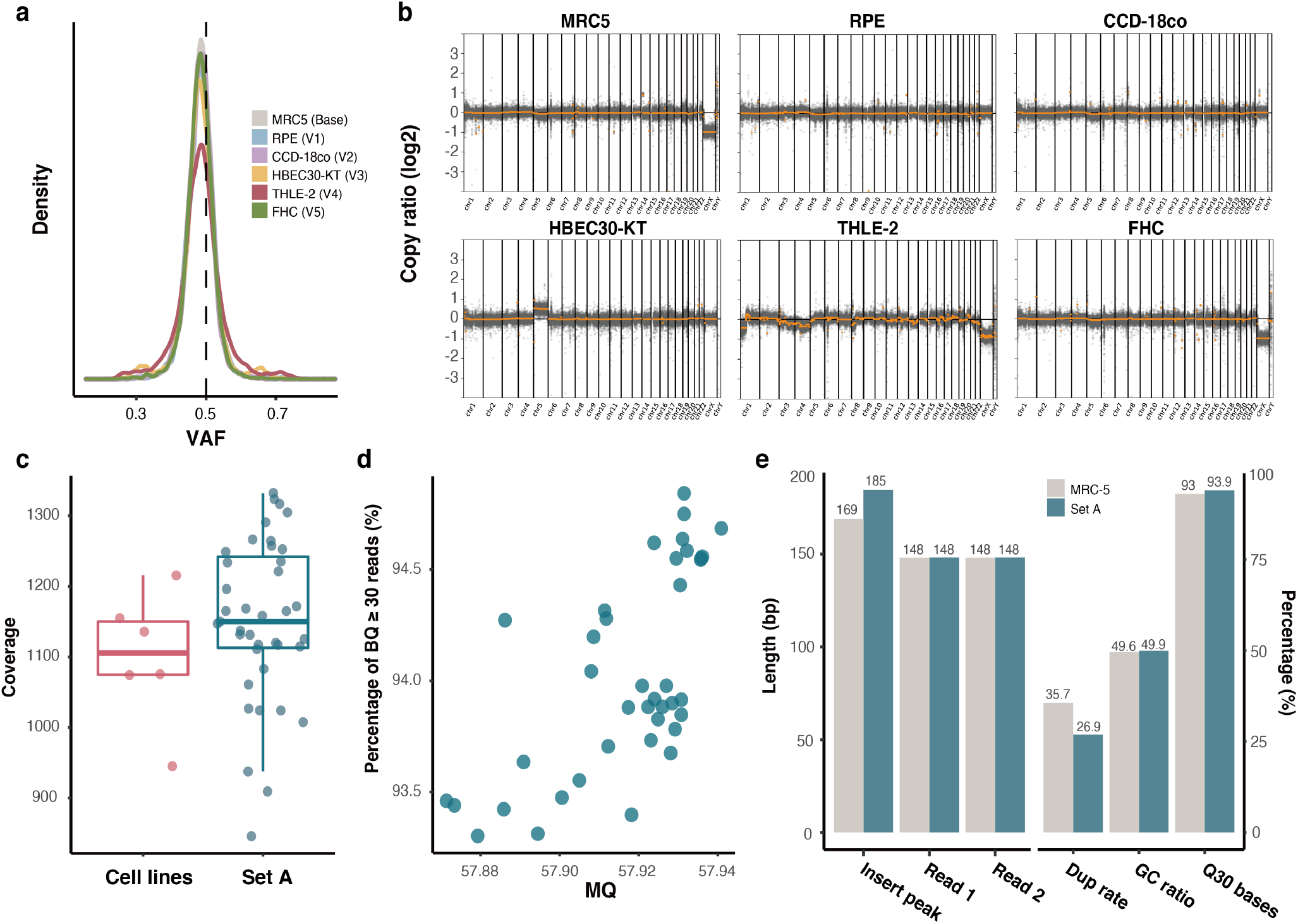
Quality validation for materials and sequencing data. **(a)** Distribution of heterozygous SNPs of the six cell lines. **(b)** Copy number ratio of the six cell lines. **(c)** Sequence coverage of the 6 cell lines and 39 mixtures of Set A. **(d)** Distribution of mean MQ and the percentage of bases with BQ over 30 in Set A. **(e)** Comparison of the read features between WES data of MRC-5 and mean values of 39 Set A. The lengths of insert size peak, paired read 1, paired read 2 and the percentages of PCR duplication rate (Dup rate), GC ratio, proportion of high-quality bases (BQ30) were compared. SNP single nucleotide polymorphism, MQ mapping quality, BQ base quality, WES whole exome sequencing

#### Sequencing quality validation

We validated 45 WES data generated in this study, including the sequencing reads of 6 cell lines and 39 mixtures. We calculated the percentage of bases with phred-scaled base quality over 30, establishing an average value of 93.93% and a minimum of 91.82% among all data. The average GC content was 49.87%, with a maximum of 51.27%, thereby depicting a very low rate of bias during library preparation. FastQC and Qualimap were also applied to validate multiple quality of sequenced reads. Sequence quality of bases in read ends had steadily high base quality over 30. Data of both cell lines and mixtures showed high coverage, with more than 1100× on average (Figure 3c). We provided WES data with high coverage and quality for cell lines as well as set A to collect reliable germline variants and remove somatic variants with high VAF, which could serve as confounding factors when selected as positive controls. The mean mapping quality and base percentage with high-quality (BQ ≥ 30) of set A are shown in Figure 3d. We also compared multiple features of reads from 39 set A and MRC5 data, which were merged when generating Set B. However, no significant differences were found, inferring that set B is less likely to have bias of two different sources (Figure 2e).

**Figure 3.**
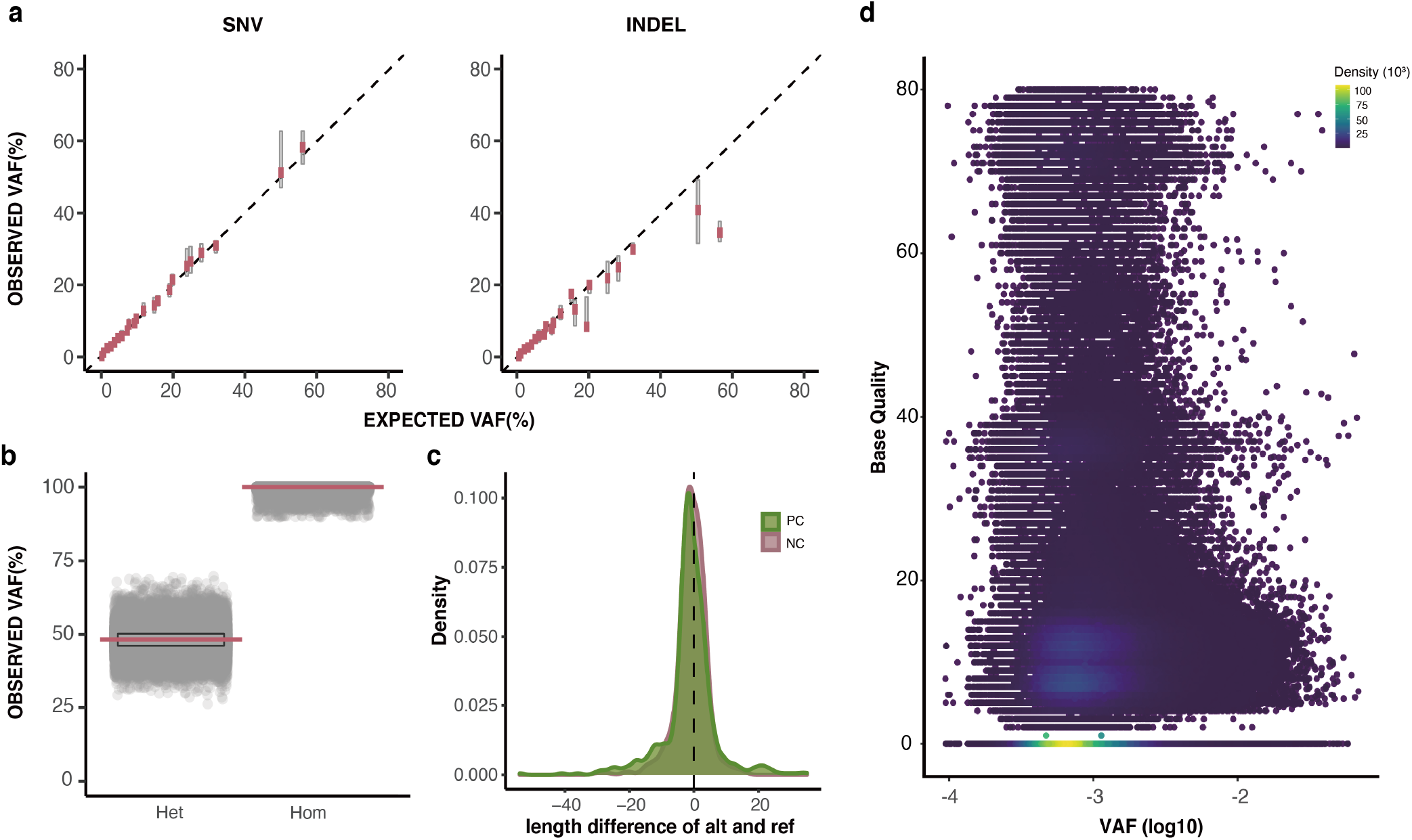
Quality validation of positive and negative controls. **(a)** Pearson correlation of expected and observed VAF of positive SNVs (r = 0.97, p < 2.2e-16) and indels (r = 0.90, p < 2.2e-16). Red lines: median VAF of observed VAFs. **(b)** VAF distributions of germline negative controls. **(c)** Length difference between alternative allele and reference allele of PC indels and NC germline indels. **(d)** Base qualities and log 10 transformed VAFs of artifacts in chromosome 1 of the random sample M2-5. The density of qualitative and quantitative distribution of artifacts were calculated from unexpected alternative alleles in Set A negative controls and depicted using ggpointdensity. VAF variant allele frequency, Het heterozygous, Hom homozygous, PC positive control, NC negative control

#### Quality Validation of positive and negative control

First, to validate the quality of positive controls, we investigated the correlations between expected VAFs of the design and observed VAFs in set A. Both SNVs and INDELs in the entire range of VAFs had a high coefficient of Pearson correlation between expected VAFs and the median value of observed VAFs among all positions with the same expected VAF (r = 0.97, p < 2.2e-16 and r = 0.91, p < 2.2e-16, respectively, Figure 3a). Thereafter, we assessed the distribution of germline negative controls in set B. The distribution of heterozygous and homozygous SNPs and INDELs is shown in Figure 3b. The length of indels in positive controls and germline negative controls demonstrated a similar distribution, indicating that they could be comparably adjusted to variant callers for performance evaluation (Figure 3c). In addition, the count of INDELs displayed a resemblance between them and most had a length smaller than 5 base pairs. Finally, we identified the quantitative and qualitative aspects of non-variant negative controls in set A. The raw alternative alleles were counted using SAMtools mplileup.

A median of 10,202,428 positions with unexpected alternative alleles was found in set A, which was approximately 30% of the total targeted regions in ultra-high depth data (1100×). Investigating the sites containing various read features would yield meaningful information. For instance, in Figure 3d, we demonstrated those sites within chromosome 1 of the randomly selected sample M2-5 with their base quality and VAFs. They had a wide range of base quality, from 0 to 80, and artifacts were concentrated at VAF near 0.001, with a base quality of zero. However, a notable number of artifacts was found with high base quality, and the destructive effect of these artifacts is assumed to be greater in data with low-sequencing coverage.

### Usage Notes

Each pair of reference standards, namely, set A and set B, can be applied to detection methods and the resultant variant calls and their properties can be assessed via a comparison to the list of positive and negative controls provided in GitHub^42^. The variant compositions as well as allele frequencies of the complete set of samples are shown in Table 2.

## Code Availability

The scripts used for constructing reference standards are available in a public repository GitHub^42^ and are accompanied by markdowns for a step-by-step description.

## Acknowledgments

This research was supported by the National Research Foundation of Korea (NRF) grant funded by the Korea government (MSIT) (No. 2019R1A2C2008050) and Korea Health Technology R&D project through the Korea Health Industry Development Institute (HI14C1324).

## Author Contributions

SK conceived the study design, prepared the manuscript.

YH developed the main analysis under the supervision of SK, prepared the manuscript.

MO conducted cell line culture and mixing under the supervision of HK.

JHK contributed to the establishment of the reference standards.

JSK and SSK performed quality validations for the properties.

All authors read and approved the final manuscript.

## Competing Interests

The authors declare no competing interests.

## Notes

### Competing Interest Statement

The authors have declared no competing interest.

## References

1 Thorpe, J., Osei-Owusu, I. A., Avigdor, B. E., Tupler, R. & Pevsner, J. Mosaicism in Human Health and Disease. Annu Rev Genet 54, 487–510, doi:10.1146/annurev-genet-041720-093403 (2020).

2 Martincorena, I. & Campbell, P. J. Somatic mutation in cancer and normal cells. Science 349, 1483–1489, doi:10.1126/science.aab4082 (2015).

3 Breuss, M. W. et al. Autism risk in offspring can be assessed through quantification of male sperm mosaicism. Nat Med 26, 143–150, doi:10.1038/s41591-019-0711-0 (2020).

4 D’Gama, A. M. & Walsh, C. A. Somatic mosaicism and neurodevelopmental disease. Nat Neurosci 21, 1504–1514, doi:10.1038/s41593-018-0257-3 (2018).

5 Freed, D. & Pevsner, J. The Contribution of Mosaic Variants to Autism Spectrum Disorder. PLoS Genet 12, e1006245, doi:10.1371/journal.pgen.1006245 (2016).

6 Lim, E. T. et al. Rates, distribution and implications of postzygotic mosaic mutations in autism spectrum disorder. Nat Neurosci 20, 1217–1224, doi:10.1038/nn.4598 (2017).

7 Rodin, R. E. et al. The landscape of somatic mutation in cerebral cortex of autistic and neurotypical individuals revealed by ultra-deep whole-genome sequencing. Nat Neurosci 24, 176–185, doi:10.1038/s41593-020-00765-6 (2021).

8 de Kock, L. et al. High-sensitivity sequencing reveals multi-organ somatic mosaicism causing DICER1 syndrome. J Med Genet 53, 43–52, doi:10.1136/jmedgenet-2015-103428 (2016).

9 Park, J. S. et al. Brain somatic mutations observed in Alzheimer’s disease associated with aging and dysregulation of tau phosphorylation. Nat Commun 10, 3090, doi:10.1038/s41467-019-11000-7 (2019).

10 Singh, S. M., Castellani, C. A. & Hill, K. A. Postzygotic Somatic Mutations in the Human Brain Expand the Threshold-Liability Model of Schizophrenia. Front Psychiatry 11, 587162, doi:10.3389/fpsyt.2020.587162 (2020).

11 Serra, E. G. et al. Somatic mosaicism and common genetic variation contribute to the risk of very-early-onset inflammatory bowel disease. Nat Commun 11, 995, doi:10.1038/s41467-019-14275-y (2020).

12 Zhu, M. et al. Somatic Mutations Increase Hepatic Clonal Fitness and Regeneration in Chronic Liver Disease. Cell 177, 608–621 e612, doi:10.1016/j.cell.2019.03.026 (2019).

13 Abyzov, A. et al. One thousand somatic SNVs per skin fibroblast cell set baseline of mosaic mutational load with patterns that suggest proliferative origin. Genome Res 27, 512–523, doi:10.1101/gr.215517.116 (2017).

14 Bae, T. et al. Different mutational rates and mechanisms in human cells at pregastrulation and neurogenesis. Science 359, 550–555, doi:10.1126/science.aan8690 (2018).

15 Ju, Y. S. et al. Somatic mutations reveal asymmetric cellular dynamics in the early human embryo. Nature 543, 714–718, doi:10.1038/nature21703 (2017).

16 Moore, L. et al. The mutational landscape of normal human endometrial epithelium. Nature 580, 640–646, doi:10.1038/s41586-020-2214-z (2020).

17 Huang, A. Y. et al. Distinctive types of postzygotic single-nucleotide mosaicisms in healthy individuals revealed by genome-wide profiling of multiple organs. PLoS Genet 14, e1007395, doi:10.1371/journal.pgen.1007395 (2018).

18 Martincorena, I. et al. Tumor evolution. High burden and pervasive positive selection of somatic mutations in normal human skin. Science 348, 880–886, doi:10.1126/science.aaa6806 (2015).

19 Manheimer, K. B. et al. Robust identification of mosaic variants in congenital heart disease. Hum Genet 137, 183–193, doi:10.1007/s00439-018-1871-6 (2018).

20 Dou, Y., Gold, H. D., Luquette, L. J. & Park, P. J. Detecting Somatic Mutations in Normal Cells. Trends Genet 34, 545–557, doi:10.1016/j.tig.2018.04.003 (2018).

21 McConnell, M. J. et al. Intersection of diverse neuronal genomes and neuropsychiatric disease: The Brain Somatic Mosaicism Network. Science 356, doi:10.1126/science.aal1641 (2017).

22 Krishnan, V. et al. Benchmarking workflows to assess performance and suitability of germline variant calling pipelines in clinical diagnostic assays. BMC Bioinformatics 22, 85, doi:10.1186/s12859-020-03934-3 (2021).

23 Cornish, A. & Guda, C. A Comparison of Variant Calling Pipelines Using Genome in a Bottle as a Reference. Biomed Res Int 2015, 456479, doi:10.1155/2015/456479 (2015).

24 Chen, Z. et al. Systematic comparison of somatic variant calling performance among different sequencing depth and mutation frequency. Sci Rep 10, 3501, doi:10.1038/s41598-020-60559-5 (2020).

25 Chen, J., Li, X., Zhong, H., Meng, Y. & Du, H. Systematic comparison of germline variant calling pipelines cross multiple next-generation sequencers. Sci Rep 9, 9345, doi:10.1038/s41598-019-45835-3 (2019).

26 Krusche, P. et al. Best practices for benchmarking germline small-variant calls in human genomes. Nat Biotechnol 37, 555–560, doi:10.1038/s41587-019-0054-x (2019).

27 Zhao, S., Agafonov, O., Azab, A., Stokowy, T. & Hovig, E. Accuracy and efficiency of germline variant calling pipelines for human genome data. Sci Rep 10, 20222, doi:10.1038/s41598-020-77218-4 (2020).

28 Zook, J. M. et al. A robust benchmark for detection of germline large deletions and insertions. Nat Biotechnol 38, 1347–1355, doi:10.1038/s41587-020-0538-8 (2020).

29 Zook, J. M. et al. Integrating human sequence data sets provides a resource of benchmark SNP and indel genotype calls. Nat Biotechnol 32, 246–251, doi:10.1038/nbt.2835 (2014).

30 Kim, J. et al. The use of technical replication for detection of low-level somatic mutations in next-generation sequencing. Nat Commun 10, 1047, doi:10.1038/s41467-019-09026-y (2019).

31 Youssoufian, H. & Pyeritz, R. E. Mechanisms and consequences of somatic mosaicism in humans. Nat Rev Genet 3, 748–758, doi:10.1038/nrg906 (2002).

32 Fernandez, L. C., Torres, M. & Real, F. X. Somatic mosaicism: on the road to cancer. Nat Rev Cancer 16, 43–55, doi:10.1038/nrc.2015.1 (2016).

33 Chen, S., Zhou, Y., Chen, Y. & Gu, J. fastp: an ultra-fast all-in-one FASTQ preprocessor. Bioinformatics 34, i884–i890, doi:10.1093/bioinformatics/bty560 (2018).

34 Li, H. & Durbin, R. Fast and accurate short read alignment with Burrows-Wheeler transform. Bioinformatics 25, 1754–1760, doi:10.1093/bioinformatics/btp324 (2009).

35 Okonechnikov, K., Conesa, A. & Garcia-Alcalde, F. Qualimap 2: advanced multi-sample quality control for high-throughput sequencing data. Bioinformatics 32, 292–294, doi:10.1093/bioinformatics/btv566 (2016).

36 Kim, S. et al. Strelka2: fast and accurate calling of germline and somatic variants. Nat Methods 15, 591–594, doi:10.1038/s41592-018-0051-x (2018).

37 Poplin, R. et al. A universal SNP and small-indel variant caller using deep neural networks. Nat Biotechnol 36, 983–987, doi:10.1038/nbt.4235 (2018).

38 Talevich, E., Shain, A. H., Botton, T. & Bastian, B. C. CNVkit: Genome-Wide Copy Number Detection and Visualization from Targeted DNA Sequencing. PLoS Comput Biol 12, e1004873, doi:10.1371/journal.pcbi.1004873 (2016).

39 Robinson, J. T. et al. Integrative genomics viewer. Nat Biotechnol 29, 24–26, doi:10.1038/nbt.1754 (2011).

40 Li, H. et al. The Sequence Alignment/Map format and SAMtools. Bioinformatics 25, 2078–2079, doi:10.1093/bioinformatics/btp352 (2009).

41 Quinlan, A. R. & Hall, I. M. BEDTools: a flexible suite of utilities for comparing genomic features. Bioinformatics 26, 841–842, doi:10.1093/bioinformatics/btq033 (2010).

42 Yoo-Jin Ha, J. K., Jisoo Kim, Sangwoo Kim* Pipeline for construction of mosaic reference standards, <https://github.com/Yonsei-TGIL/Mosaic-Reference-Standards.git> (2021).

43 Yoo-Jin Ha, J. K., Jisoo Kim, Sangwoo Kim* Yonsei-TGIL/Mosaic-Reference-Standards: (v1.0.1). Zenodo, doi:https://doi.org/10.5281/zenodo.5338953 (2021).

44 Ramirez, R. D. et al. Immortalization of human bronchial epithelial cells in the absence of viral oncoproteins. Cancer Res 64, 9027–9034, doi:10.1158/0008-5472.CAN-04-3703 (2004).

